# stormTB: A web-based simulator of a murine minimal-PBPK model for anti-tuberculosis treatments

**DOI:** 10.1101/2024.07.04.602069

**Authors:** Roberto Visintainer, Anna Fochesato, Daniele Boaretti, Stefano Giampiccolo, Shayne Watson, Micha Levi, Federico Reali, Luca Marchetti

## Abstract

**Introduction:** Tuberculosis (TB) poses a significant threat to global health, with millions of new infections and approximately one million deaths annually. Various modeling efforts have emerged, offering tailored data-driven and physiologically-based solutions for novel and historical compounds. However, this diverse modeling panorama may lack consistency, limiting result comparability. Drug-specific models are often tied to commercial software and developed on various platforms and languages, potentially hindering access and complicating the comparison of different compounds.

**Methods:** This work introduces stormTB: SimulaTOr of a muRine Minimal-pbpk model for anti-TB drugs. It is a web-based interface for our minimal physiologically based pharmacokinetic (mPBPK) platform, designed to simulate custom treatment scenarios for tuberculosis in murine models. The app facilitates visual comparisons of pharmacokinetic profiles, aiding in assessing drug-dose combinations.

**Results:** The mPBPK model, supporting 11 anti-TB drugs, offers a unified perspective, overcoming the potential inconsistencies arising from diverse modeling efforts. The app, publicly accessible, provides a user-friendly environment for researchers to conduct what-if analyses and contribute to collective TB eradication efforts. The tool generates comprehensive visualizations of drug concentration profiles and pharmacokinetic/pharmacodynamic indices for TB-relevant tissues, empowering researchers in the quest for more effective TB treatments. stormTB is freely available at the link: https://apps.cosbi.eu/stormTB.

## Introduction

Tuberculosis (TB) poses a significant threat to global health, with millions of new infections and approximately 1.3 million deaths annually (World Health Organization, 2023). Recent initiatives have spurred research in the field, with several novel drug candidates rekindling a pipeline that was nearly empty a decade ago (Dartois & Rubin, 2022). In this context, computational models play a crucial role by rapidly providing information on the exposure and efficacy of new compounds, expediting the drug development process, and aiding in prioritizing the most promising candidates (Dartois & Rubin, 2022).

Alongside this renewed momentum in TB research, various modeling efforts have emerged, offering tailored solutions for both novel and historical compounds. These models suggest dosing strategies and elucidate the effectiveness of monotherapies and drug combinations, *i*.*e*., regimens. Data-driven solutions, such as pharmacokinetic (PK) and pharmacokinetic/pharmacodynamic (PK/PD) models, provide effective means to derive exposure and efficacy indexes for TB compounds (Alffenaar et al., 2022; Ernest et al., 2023; Wicha et al., 2018; Zhang et al., 2020). Simultaneously, physiologically based approaches, including PBPK and semi- or fully mechanistic models, provide insights into the intricate diffusion of anti-TB compounds within TB lesions, revealing scenarios that may differ among drugs (Ernest et al., 2021; Humphries et al., 2021; Mehta et al., 2023; Muliaditan & Della Pasqua, 2022).

While the rich and varied landscape of modeling efforts presents valuable insights, it sometimes faces challenges in maintaining uniformity, which can affect the comparability of results. The creation of drug-specific models frequently relies on proprietary software and unfolds across diverse platforms and programming languages, potentially hindering access and complicating the comparison of different compounds. A recent approach is to rely on web interfaces that generate, tune, and simulate (PB)PK models for a wide range of applications. Some examples include igPBPK, an R-based Shiny app for simulating drug withdrawal intervals in cattle or swine for flunixin, florfenicol, and penicillin G with a PBPK model (Chou et al., 2022). ModVizPop is another web app to simulate PK/PD dynamics for compartmental modeling (Vaddady & Kandala, 2021). E-campsis and gPKPDviz are freemium R Shiny apps developed by Calvagone and Genentech, respectively, that allow simulating the PK/PD dynamics with a collection of PK models (Lu et al., 2024; Luyckx, 2024) and the custom integration of thresholds and AUC to be displayed in the plots. Still, there is a need for an open-source unified tool specific to anti-TB PBPK drug analysis.

Here, we present a web-based tool tailored for anti-TB drug dynamics and PK/PD metrics in the treatment scenarios to support model-informed treatment development under a unified perspective. It leverages a previously published minimal physiologically based pharmacokinetic (mPBPK) model that supports 11 historical and under-development anti-TB drugs in a common pre-clinical murine model (Reali et al., 2024). We have then consolidated the results from model calibration and virtual population-based uncertainty quantification in the herein introduced R-based web app, stormTB, streamlining model inspection and enabling users to conduct independent what-if analyses using the mPBPK platform. Users can compose a treating scenario by selecting one drug from the 11 originally included in (Reali et al., 2024), the dosage and the treatment length. Scientists can iteratively adjust the experimental settings based on the simulated PK/PD performance and save the resulting scenario in the workspace. Up to four precomputed monotherapy scenarios can be selected from the workspace and visualized in combination for comparative analysis.

In addition to mean PK profiles, an option for generating a virtual population (VP) is available, offering suitable choices for population size and the coefficient of variation governing the sampling of clearance and absorption rate values in the population. The tool produces comprehensive visualizations of the drug concentration profile in all nine compartments comprising the mPBPK model, along with descriptions of the PK/PD indices for TB-relevant tissues.

stormTB is freely available at the link: https://apps.cosbi.eu/stormTB

## Methods

### Minimal-PBPK model

stormTB implements the minimal physiologically based pharmacokinetics model (mPBPK) presented by (Reali et al., 2024) that describes the disposition of 11 anti-pulmonary-TB drugs in murine models. The supported drugs are rifampicin (RIF, R), rifapentine (RPT, P), pyrazinamide (PZA, Z), ethambutol (EMB, E), isoniazid (INH, H), moxifloxacin (MOX, M), delamanid (DEL), pretomanid (PRE, Pa), bedaquiline (BDQ, B), Quabodepistat (QBS, OPC-167832), and GSK2556286 (G286).

The mPBPK model consists of nine ordinary differential equations obtained by streamlining a whole-body mPBPK model via the identification of the tissues least involved in the TB site of action, and the combination of relative compartments to obtain a smaller set of equations (Nestorov et al., 1998; Ryu et al., 2022). Out of the 25 model parameters, only the absorption rate (Ka) and the total clearance (CL) were calibrated using mouse data for each of the 11 drugs and are presented in the app description. A complete list of model parameters is available in (Reali et al., 2024).

### Implementation and simulations

The original model (Reali et al., 2024), implemented in MATLAB, has been translated into R (4.3.1) and C (gcc 11.4.0) to reduce simulation time and executed using the deSolve (1.40) package (Soetaert et al., 2010). The stormTB user interface is developed with Shiny (1.7.5.1), shinyBS (0.61.1), shinyhelper (0.3.2), shinycssloaders (1.0.0), shinyWidgets (0.8.0), shinyjs (2.1.0), dplyr (1.1.3), tidyr (1.3.0), collapse (2.0.13), zip (2.3.0), ggplot2 (3.4.4), scales (1.2.1), ggiraph(0.8.7) (Attali, 2021; Bailey, 2022; Chang et al., 2024; Csárdi et al., 2023; Gohel & Skintzos, 2023; Krantz, 2024; Mason-Thom, 2019; Perrier et al., 2024; Sali & Attali, 2020; Wickham, 2016; Wickham, François, et al., 2023; Wickham, Pedersen, et al., 2023; Wickham, Vaughan, et al., 2023).

Each simulation can be computed on a single mouse, applying the reference clearance and absorption rates, or on a virtual population of mice. In the latter case, the user can specify the Number of Mice (default 100) to be simulated, along with the coefficients of variation of the sampled model parameters (CV clearance and CV absorption, both defaulting to 0.2). These values are applied via a lognormal distribution to the two parameters of the specified drug. With the VP option activated, all the statistics are presented with their confidence intervals computed as 5 – 95% of the distributions of the values. The same CIs are represented in all the plots.

To generate the dynamics of virtual populations, stormTB particularly benefits from the translation of the model in C, being able to simulate a test case with a virtual population of 1000 mice (RIF, 15 days, and default parameters) in 4.3 seconds (Table 1). This is in clear contrast to the 20 minutes required using a pure R implementation, resulting in a computational time decrease of more than 287-fold. Moreover, for fast postprocessing, we implemented data transformation and basic statistical computation using the R library collapse (written in C/C++). This solution reduced to just 9 seconds the time needed to compute the test case from the start of the simulation to the rendering of the resulting plots and tables.

**Table 1:** Comparison between pure R and C implementations of computational time for the pure simulation process. We consider 6 different scenarios: number of mice in the virtual population (N VP) [500, 100] and number of doses (N Doses) [5, 10, 15]. We report the median and standard deviation measurement obtained with 5 repetitions of a simulation of rifampicin and default parameters.

| N VP |  | 500 |  |  | 1000 |  |  |
| --- | --- | --- | --- | --- | --- | --- | --- |
| N Doses |  | 5 | 10 | 15 | 5 | 10 | 15 |
| Time (sec)<br>± SD | R | **195.45**<br>±2.061 | **385.26**<br>±3.185 | **603**<br>±7.901 | **393.54**<br>±3.751 | **801**<br>±6.125 | **1244.4**<br>±24.8 |
|  | C | **1.155**<br>±0.087 | **1.667**<br>±0.045 | **2.23**<br>±0.062 | **2.347**<br>±0.066 | **3.304**<br>±0.124 | **4.335**<br>±0.129 |
| Fold Change |  | 169.7 | 231.2 | 270.4 | 167.7 | 242.5 | 287.1 |

### Options

stormTB provides a platform to simulate custom anti-pulmonary-TB treatment scenarios in which the user can choose between the 11 supported drugs and can select the dose amount expressed in mg/kg and number of doses to be administered. When a specific drug is selected from the dropdown menu, the dose is automatically set to its human equivalent as a reference value. Users can then freely adjust this value as needed. The number of doses also sets the duration of the simulation since the model assumes the most common setup for anti-TB treatments, which is one oral administration per day. Once the simulation is completed, the user obtains plots and tables summarizing the predictions about the pharmacokinetics (PK) and pharmacodynamics (PD) of the defined scenario.

The default PK plots showcase the drug concentration profile in plasma and lung compartments (Figure 1), alternatively the users have the option to analyze PK profiles for all nine model compartments as shown in Figure 2. To better appreciate the dynamics, the visualization can be switched to the logarithmic scale with an adjustable lower limit on the y axis. For plasma and lungs, the plots can include the values of the minimal inhibitory concentration at which at least 50% of the isolates in a test population are inhibited (MIC), the minimal bactericidal concentration at which at least 90% of the isolates are killed (MBC), the concentration that inhibits 90% of growth in macrophages (MacIC) and the Wayne cidal concentration or concentration that kills 90% of extracellular M. tuberculosis under anaerobic conditions (WCC) (Lakshminarayana et al., 2015; Wayne & Hayes, 1996). Furthermore, these plots are complemented by tissue-specific tables summarizing important PK and PD indices from the simulation, *i*.*e*., the maximum achieved concentration (Cmax), the time at which the maximum concentration is reached (Tmax), the area under the curve for the total amount of drug and for the fraction unbound (AUC, fuAUC), and the time above the potency thresholds (T > MIC50, T > MBC90, T > MacIC90 and T > WCC90). The times above the potency thresholds are computed considering the 24 hours after the last simulated dose. The AUC is calculated on a configurable interval between 1 and the end of the simulation, see Figure 3.

**Figure 1:**
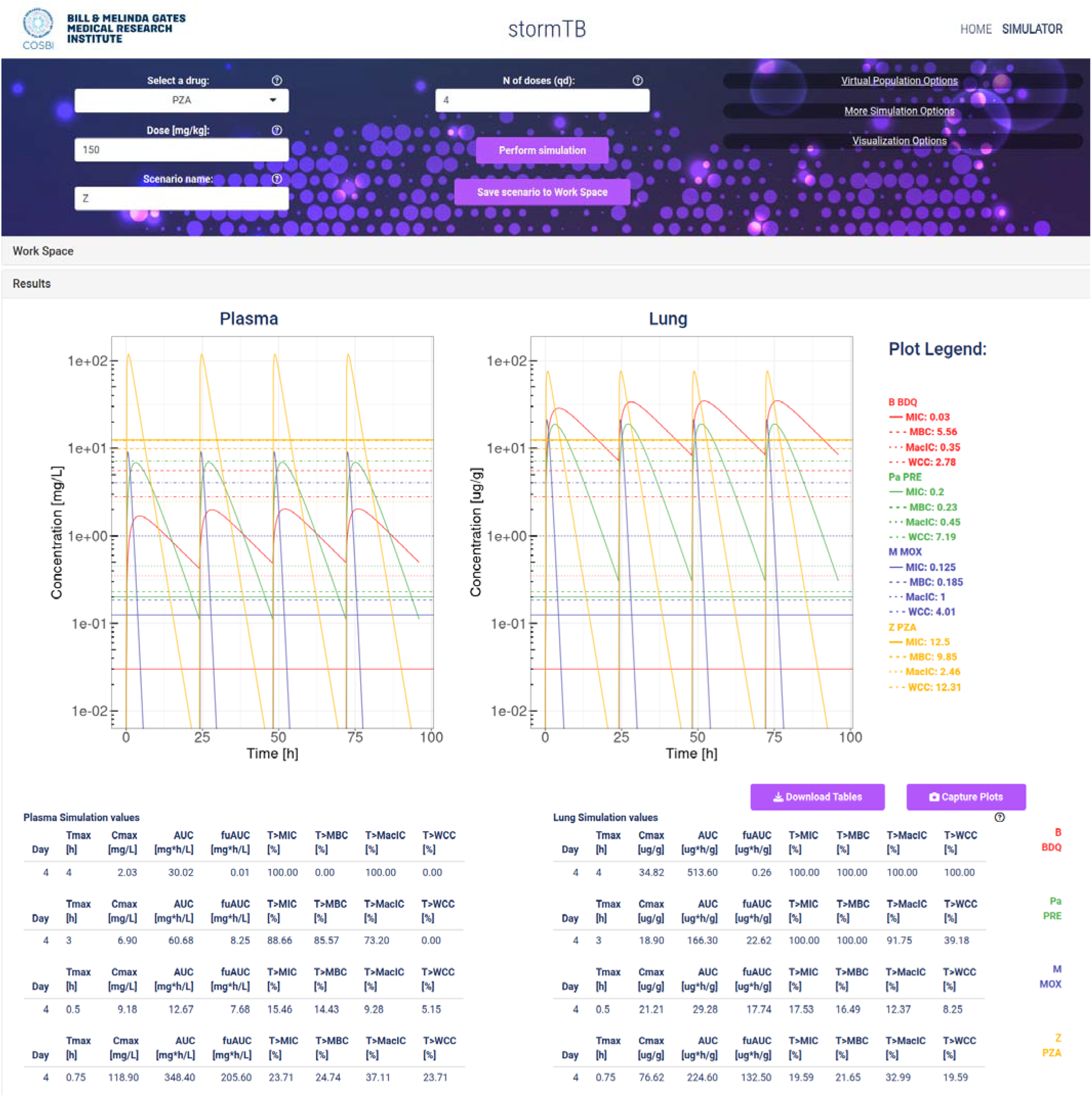
An example of simulation considering the TB drugs involved in the BPaMZ regimen: 25 mg/kg of bedaquiline (B), 10 mg/kg of pretomanid (Pa), 100 mg/kg of moxifloxacin (M), and 150 mg/kg of pyrazinamide (Z). The top part of the figure shows the input panel. The central part provides the predicted dynamics of the drug concentrations in plasma (mg/L) and lung (ug/g). Additionally, it shows the drug values for the minimal inhibition concentration (MIC), the minimum bactericidal concentration (MBC), the minimal inhibition concentration in macrophages (MacIC), and the Wayne cidal concentration (WCC). Note that lung density is assumed 1 g/ml, allowing for direct concentration comparison. The bottom of the figure reports the simulation statistics and refers to the last day of the simulated study.

**Figure 2:**
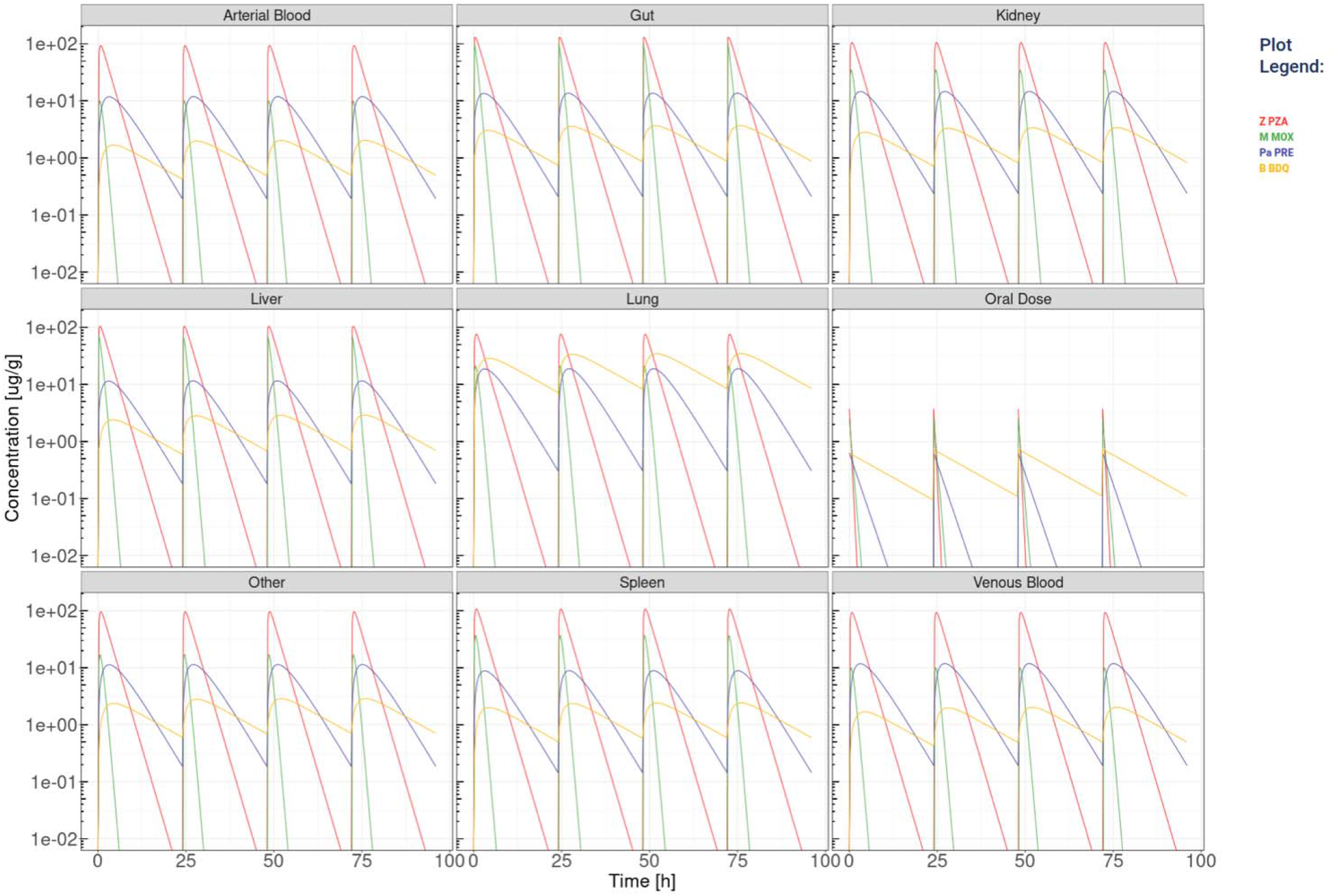
The visualization produced for all the compartments’ simulation considering the TB drugs involved in the BPaMZ regimen: 25 mg/kg of bedaquiline (B), 10 mg/kg of pretomanid (Pa), 100 mg/kg of moxifloxacin (M), and 150 mg/kg of pyrazinamide (Z). Here the logarithmic visualization is applied to better appreciate the different dynamics of the drugs.

**Figure 3:**
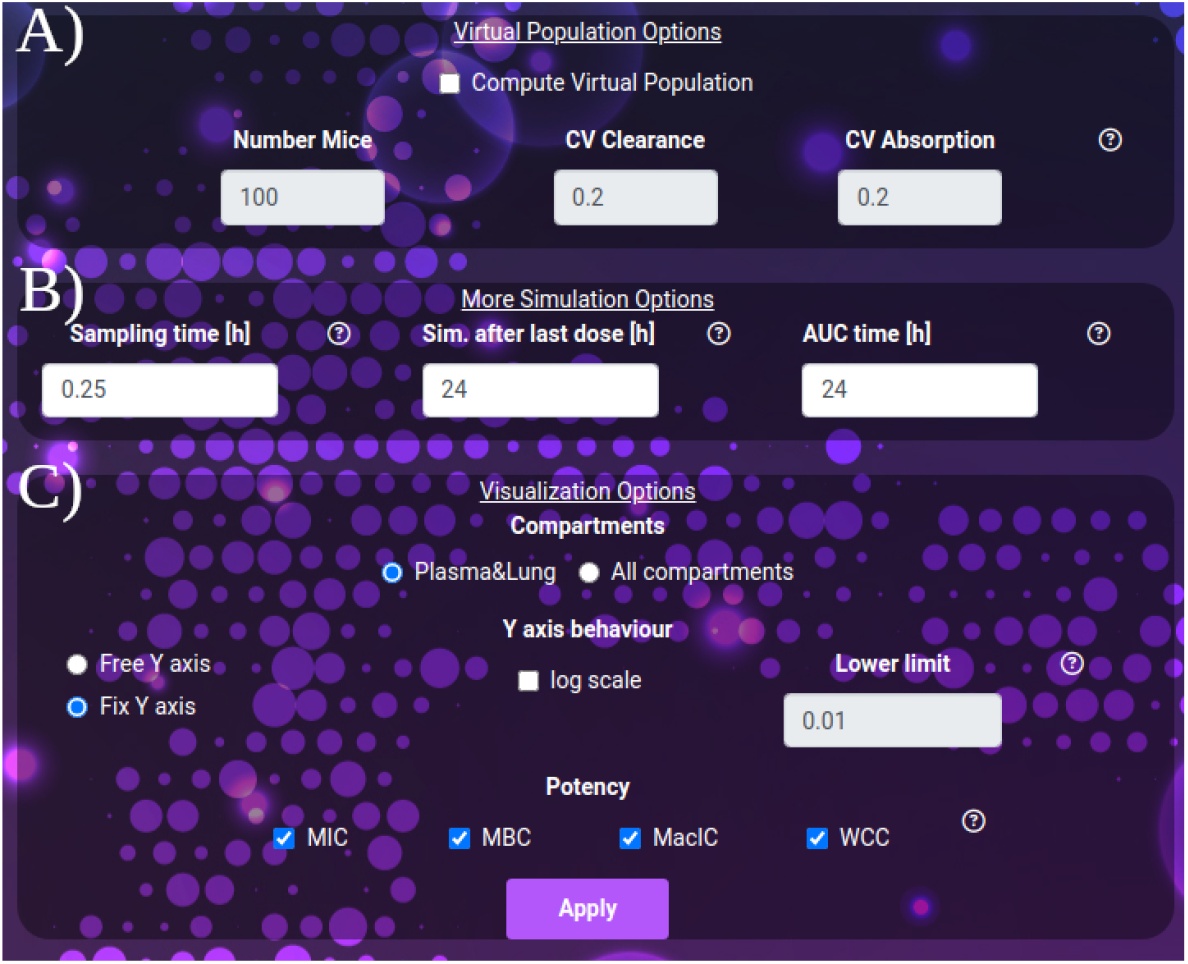
A) The virtual population options: number of simulated mice and coefficients of variation to apply to clearance and absorption. B) The extra simulation parameters: the sampling time, the interval to be simulated after the last dose and the interval to be considered for the computation of the AUC after the last dose. C) The visualization parameters: selection between the plasma-lung visualization and all compartments. The y-axis behavior: free or fixed scale, application of logarithm and setting of the lower limit of the scale. The potency section: selection of the thresholds to be considered in the plot and result tables.

For both single mouse and virtual populations (Figures 1, 2, and 4), the web-app offers the option to store and recall simulated scenarios in the workspace area. When recalling saved simulations, the user can visually compare PK profiles of different scenarios, analyzing the impact of various drug-dose combinations with the aid of a combined plot and all the statistics (Figures 1 and 2). To guarantee a tidy visualization, the comparison tool supports a maximum of four scenarios of the same length. Plots, tables and parameters of the scenarios can be easily downloaded for reporting purposes, moreover, the simulated data can be saved in raw format for successive analysis with external tools and possibly integrated with results from other software.

**Figure 4:**
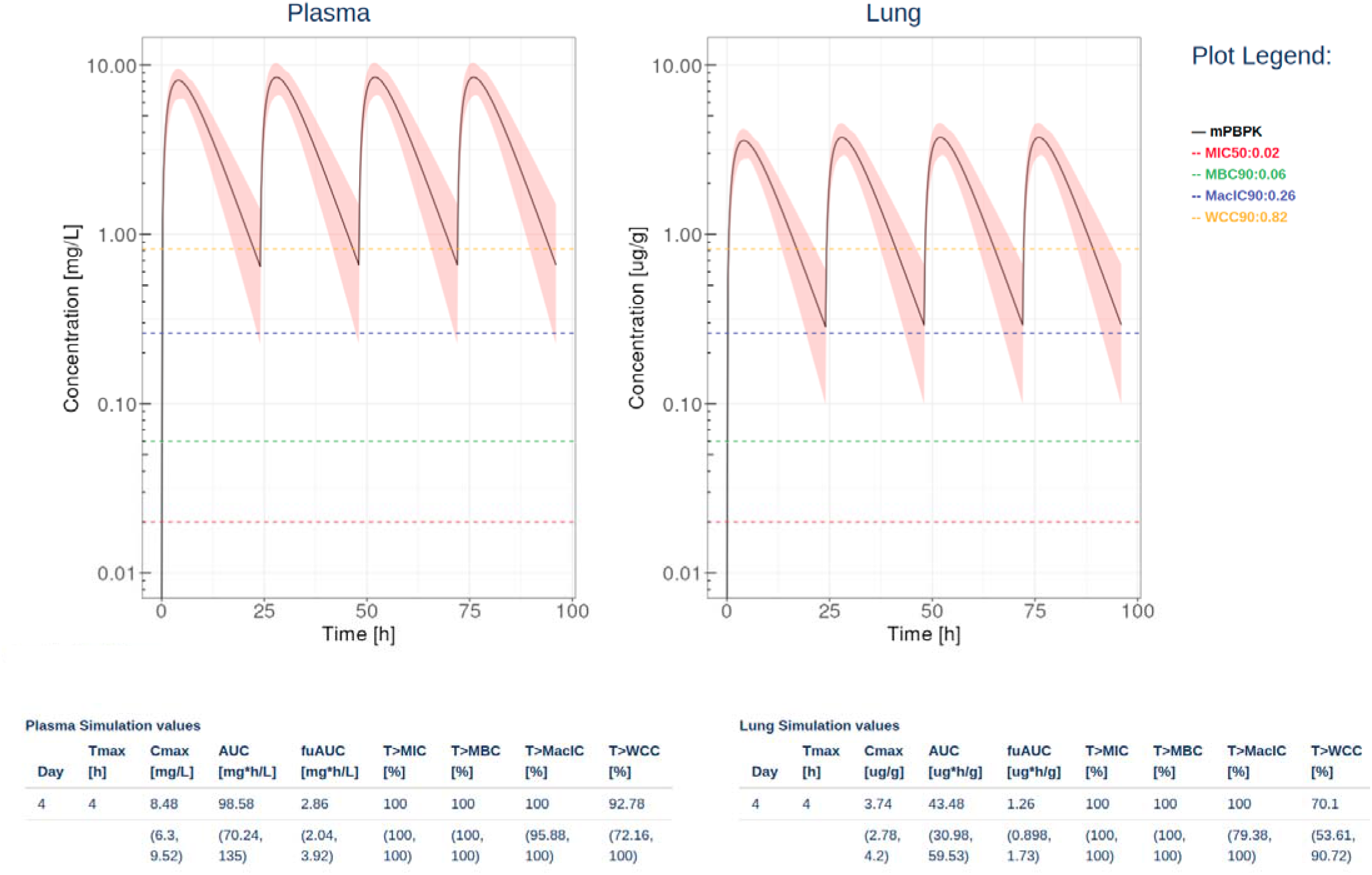
An example of simulation computed with Rifampicin at a dose of 10mg/kg, 4 doses and a virtual population of 100 mice and coefficient of variation so 0.2 for both clearance and absorption. The tables report the computed statistics showing the last day mean and 5-95 quantiles of the virtual mice.

#### Visualization options include

- The simulation Sampling time, expressed in hours (default 0.25, min: 0.1, max: 1), governs the density of simulated points. It affects the post-process time to render the images.
- The duration of the simulation after the last dose, expressed in hours (default and minimum value 24 hours, max: 1000). Increasing this parameter allows the user to exhaustively analyze the dynamics of drugs with very long clearance time such as bedaquiline or rifapentine.
- AUC time, expressed in hours (default 24), indicates the number of hours after the last dose, for which the AUC is computed, therefore AUC time must be equal or smaller than simulation after the last dose. AUC time does not apply to the computation of the time above the potency thresholds, which always refers to 24 hours after the last dose.
- For the Y axis, the following options are available: activate the logarithmic scale and set a Y axis lower limit, set the same limit for all the plots shown (Fix Y axis), or automatically adjust to the compartment dynamics (Free Y axis).

## Results

We present stormTB: SimulaTOr of a muRine Minimal-pbpk model for anti-TB drugs, a web-based tool to interact with our minimal PBPK platform to simulate treatment scenarios for tuberculosis in mice. Pharmacokinetics results are reported in terms of AUC, Cmax, and Tmax. In addition, the user is presented with four pharmacodynamics measures expressed as the percentage of the treatment time above each threshold (MIC, MBC, MacIC, and WCC). The use of a customizable virtual population allows the user to enrich the results with confidence intervals for each quantitative result. To enhance the interoperability of stormTB, we made the raw simulated data available ensuring seamless integration and enhanced flexibility for users to add the results to their projects.

An analysis using our web app typically begins with defining the scenarios under investigation, each representing a single drug treatment with a dose administered once a day and a specified duration for the simulated experiment. In this section, we present an example of comparison considering the drugs constituting the BPaMZ regimen: 25 mg/kg of bedaquiline (B), 10 mg/kg of pretomanid (Pa), 100 mg/kg of moxifloxacin (M), and 150 mg/kg of pyrazinamide (Z).

The comparative assessment across simulated monotherapy treatments and tissues leads to insights into their effectiveness (Figure 1). Specifically, pyrazinamide is the only component of the regimen less abundant in the lung than in plasma, reaching the highest penetration among those under investigation in terms of Cmax. However, it shows poor performance in terms of time above MIC and MBC in the lung during the last day of treatment, with 19.59% and 21.65%, respectively. The simulations showed that bedaquiline, pretomanid, and moxifloxacin accumulate more in the lung than in plasma.

MIC and MBC thresholds are reached for 100% of the treatment by bedaquiline and pretomanid, while moxifloxacin, on the last day of treatment, is above the MIC and MBC for 17.53% and 16.49% of the time, respectively. Moreover, bedaquiline performs best in terms of WCC and MacIC, reaching both thresholds throughout the whole treatment. The second-best performing drug is again pretomanid, with 91.75% and 39.18%, respectively, showing the importance of these two compounds in the regimen to reduce the hard-to-treat bacterial and be effective in TB-lesions (Dartois & Rubin, 2022; Reali et al., 2024). Additionally, stormTB allows to inspect the predicted exposure in all the compartments (Figure 2).

The insights gained from the comparative analysis of the BPaMZ regimen highlight the potential of stormTB in optimizing drug combinations and dosages for enhanced therapeutic efficacy. Such analyses are instrumental for researchers in comparing various drug combinations and doses, designing novel regimens, and fine-tuning dosages to meet effective thresholds.

## Discussion

The mPBPK model simulator stormTB is a versatile, web-based application that streamlines the efforts of both modeling and non-modeling scientists in extracting crucial pharmacokinetic and pharmacodynamic measurements for supported anti-tuberculosis compounds. This simulation tool enables users to access specific PK and PD metrics, analyze compartment-specific PK profiles, and compare them with existing data, thereby promoting efficient benchmarking. While other web interfaces have been available providing a valuable support for researchers in a broad set of applications related to PBPK modeling (Chou et al., 2022; Lu et al., 2024; Luyckx, 2024; Vaddady & Kandala, 2021), we focus on TB-specific ADME dynamics and PK/PD metrics in the treatment scenarios.

Future updates to stormTB include the integration of a drug-drug interaction module, which will enable researchers to evaluate the intricate ADME dynamics of co-administered drugs, thereby shifting the focus towards the complexities of realistic treatment regimens. The expansion to incorporate additional animal models, such as rabbits for efficacy benchmarks and dogs and rats for toxicological evaluations will enhance the tool’s utility across various research domains. Supporting cross-species translations and predicting human exposure and efficacy will further bridge the gap between preclinical findings and clinical outcomes, potentially informing global health policies and refining TB treatment protocols.

As a user-friendly and freely accessible resource, stormTB democratizes the PBPK and PD analysis of a broad spectrum of drugs, both historical and novel. It empowers TB researchers globally to compare and benchmark drug combinations and dosages, thereby accelerating the discovery and optimization of treatment strategies removing the need for licenses or subscription plans. Through its contributions, stormTB aligns with the collective effort to eradicate TB by the end of the decade, aspiring to make a significant impact on public health.

## Code Availability

stormTB is freely available at https://apps.cosbi.eu/stormTB/

## References

Alffenaar, J. W. C., de Steenwinkel, J. E. M., Diacon, A. H., Simonsson, U. S. H., Srivastava, S., & Wicha, S. G. (2022). Pharmacokinetics and pharmacodynamics of anti-tuberculosis drugs: An evaluation of in vitro, in vivo methodologies and human studies. Frontiers in Pharmacology, 13, 1063453. 10.3389/FPHAR.2022.1063453/BIBTEX

Attali, D. (2021). shinyjs: Easily Improve the User Experience of Your Shiny Apps in Seconds. https://CRAN.R-project.org/package=shinyjs

Bailey, E. (2022). shinyBS: Twitter Bootstrap Components for Shiny. https://CRAN.R-project.org/package=shinyBS

Chang, W., Cheng, J., Allaire, J. J., Sievert, C., Schloerke, B., Xie, Y., Allen, J., McPherson, J., Dipert, A., & Borges, B. (2024). shiny: Web Application Framework for R. https://shiny.posit.co/

Chou, W. C., Tell, L. A., Baynes, R. E., Davis, J. L., Maunsell, F. P., Riviere, J. E., & Lin, Z. (2022). An Interactive Generic Physiologically Based Pharmacokinetic (igPBPK) Modeling Platform to Predict Drug Withdrawal Intervals in Cattle and Swine: A Case Study on Flunixin, Florfenicol, and Penicillin G. Toxicological Sciences, 188(2), 180–197. 10.1093/TOXSCI/KFAC056

Csárdi, G., Podgórski, K., & Geldreich, R. (2023). zip: Cross-Platform “zip” Compression. https://CRAN.R-project.org/package=zip

Dartois, V. A., & Rubin, E. J. (2022). Anti-tuberculosis treatment strategies and drug development: challenges and priorities. Nature Reviews Microbiology 2022 20:11, 20(11), 685–701. 10.1038/s41579-022-00731-y

Ernest, J. P., Goh, J. J. N., Strydom, N., Wang, Q., van Wijk, R. C., Zhang, N., Deitchman, A., Nuermberger, E., & Savic, R. M. (2023). Translational predictions of phase 2a first-in-patient efficacy studies for antituberculosis drugs. European Respiratory Journal, 62(2), 2300165. 10.1183/13993003.00165-2023

Ernest, J. P., Sarathy, J., Wang, N., Kaya, F., Zimmerman, M. D., Strydom, N., Wang, H., Xie, M., Gengenbacher, M., Via, L. E., Barry, C. E., Carter, C. L., Savic, R. M., & Dartois, V. (2021). Lesion Penetration and Activity Limit the Utility of Second-Line Injectable Agents in Pulmonary Tuberculosis. Antimicrobial Agents and Chemotherapy, 65(10). 10.1128/AAC.00506-21

Gohel, D., & Skintzos, P. (2023). ggiraph: Make “ggplot2” Graphics Interactive. https://CRAN.R-project.org/package=ggiraph

Humphries, H., Almond, L., Berg, A., Gardner, I., Hatley, O., Pan, X., Small, B., Zhang, M., Jamei, M., & Romero, K. (2021). Development of physiologically□based pharmacokinetic models for standard of care and newer tuberculosis drugs. CPT: Pharmacometrics & Systems Pharmacology, 10(11), 1382–1395. 10.1002/psp4.12707

Krantz, S. (2024). collapse: Advanced and Fast Statistical Computing and Data Transformation in R. https://sebkrantz.github.io/collapse/

Lakshminarayana, S. B., Huat, T. B., Ho, P. C., Manjunatha, U. H., Dartois, V., Dick, T., & Rao, S. P. S. (2015). Comprehensive physicochemical, pharmacokinetic and activity profiling of anti-TB agents. Journal of Antimicrobial Chemotherapy, 70(3), 857–867. 10.1093/jac/dku457

Lu, T., Poon, V., Brooks, L., Velasquez, E., Anderson, E., Baron, K., Jin, J. Y., & Kågedal, M. (2024). gPKPDviz: A flexible R shiny tool for pharmacokinetic/pharmacodynamic simulations using mrgsolve. CPT: Pharmacometrics & Systems Pharmacology, 13(3), 341–358. 10.1002/PSP4.13096

Luyckx, N. (2024). e-Campsis: Shiny dashboard interface for Campsis. https://github.com/Calvagone/campsis

Mason-Thom, C. (2019). shinyhelper: Easily Add Markdown Help Files to “shiny” App Elements. https://CRAN.R-project.org/package=shinyhelper

Mehta, K., Guo, T., van der Graaf, P. H., & van Hasselt, J. G. C. (2023). Predictions of Bedaquiline and Pretomanid Target Attainment in Lung Lesions of Tuberculosis Patients using Translational Minimal Physiologically Based Pharmacokinetic Modeling. Clinical Pharmacokinetics, 62(3), 519–532. 10.1007/s40262-023-01217-7

Muliaditan, M., & Della Pasqua, O. (2022). Bacterial growth dynamics and pharmacokinetic– pharmacodynamic relationships of rifampicin and bedaquiline in BALB/c mice. British Journal of Pharmacology, 179(6), 1251–1263. 10.1111/BPH.15688

Nestorov, I. A., Aarons, L. J., Arundel, P. A., & Rowland, M. (1998). Lumping of whole-body physiologically based pharmacokinetic models. Journal of Pharmacokinetics and Biopharmaceutics, 26(1), 21–46. 10.1023/a:1023272707390

Perrier, V., Meyer, F., & Granjon, D. (2024). shinyWidgets: Custom Inputs Widgets for Shiny. https://CRAN.R-project.org/package=shinyWidgets

Reali, F., Fochesato, A., Kaddi, C., Visintainer, R., Watson, S., Levi, M., Dartois, V., Azer, K., & Marchetti, L. (2024). A minimal PBPK model to accelerate preclinical development of drugs against tuberculosis. Frontiers in Pharmacology, 14. 10.3389/fphar.2023.1272091

Ryu, H. J., Kang, W. H., Kim, T., Kim, J. K., Shin, K. H., Chae, J. W., & Yun, H. Y. (2022). A compatibility evaluation between the physiologically based pharmacokinetic (PBPK) model and the compartmental PK model using the lumping method with real cases. Frontiers in Pharmacology, 13, 3057. 10.3389/fphar.2022.964049

Sali, A., & Attali, D. (2020). shinycssloaders: Add Loading Animations to a “shiny” Output While It’s Recalculating. https://CRAN.R-project.org/package=shinycssloaders

Soetaert, K., Petzoldt, T., & Setzer, R. W. (2010). Solving Differential Equations in R□: Package deSolve. Journal of Statistical Software, 33(9). 10.18637/jss.v033.i09

Vaddady, P., & Kandala, B. (2021). ModVizPop: A shiny interface for empowering teams to perform interactive pharmacokinetic/pharmacodynamic simulations. CPT: Pharmacometrics & Systems Pharmacology, 10(11), 1323–1331. 10.1002/PSP4.12697

Wayne, L. G., & Hayes, L. G. (1996). An in vitro model for sequential study of shiftdown of Mycobacterium tuberculosis through two stages of nonreplicating persistence. Infection and Immunity, 64(6), 2062–2069. 10.1128/iai.64.6.2062-2069.1996

Wicha, S. G., Clewe, O., Svensson, R. J., Gillespie, S. H., Hu, Y., Coates, A. R. M., & Simonsson, U. S. H. (2018). Forecasting Clinical Dose–Response From Preclinical Studies in Tuberculosis Research: Translational Predictions With Rifampicin. Clinical Pharmacology and Therapeutics, 104(6), 1208–1218. 10.1002/cpt.1102

Wickham, H. (2016). ggplot2: Elegant Graphics for Data Analysis. Springer-Verlag New York. https://ggplot2.tidyverse.org

Wickham, H., François, R., Henry, L., Müller, K., & Vaughan, D. (2023). dplyr: A Grammar of Data Manipulation. https://CRAN.R-project.org/package=dplyr

Wickham, H., Pedersen, T. L., & Seidel, D. (2023). scales: Scale Functions for Visualization. https://CRAN.R-project.org/package=scales

Wickham, H., Vaughan, D., & Girlich, M. (2023). tidyr: Tidy Messy Data. https://CRAN.R-project.org/package=tidyr

World Health Organization. (2023). Global tuberculosis report 2023. https://iris.who.int/.

Zhang, N., Strydom, N., Tyagi, S., Soni, H., Tasneen, R., Nuermberger, E. L., & Savic, R. M. (2020). Mechanistic Modeling of Mycobacterium tuberculosis Infection in Murine Models for Drug and Vaccine Efficacy Studies. Antimicrobial Agents and Chemotherapy, 64(3). 10.1128/AAC.01727-19

